# Transcriptome features of striated muscle aging and predictability of protein level changes

**DOI:** 10.1101/2021.06.12.448203

**Authors:** Yu Han, Lauren Z. Li, Nikhitha L. Kastury, Cody T Thomas, Maggie P. Y. Lam, Edward Lau

## Abstract

RNA and protein levels correlate only partially and some transcripts are better correlated with their protein counterparts than others. This suggests that in aging and disease studies, some transcriptomics markers may carry more information in predicting protein-level changes. Here we applied a computational data analysis workflow to predict which transcriptomic changes are more likely relevant to protein-level regulation in striated muscle aging. The protein predictability of each transcript is estimated from existing large proteogenomics data sets, then transferred to new total RNA sequencing data comparing skeletal muscle and cardiac muscle in young adult (~4 months) mice vs. early aging (~20 months) mice. Aging cardiac and skeletal muscles both invoke transcriptomic changes in innate immune system and mitochondria pathways but diverge in extracellular matrix processes. On an individual gene level, we identified 611 age-associated signatures in skeletal and cardiac muscles at 10% FDR, including a number of myokine and cardiokine encoding genes. We estimate that about 48% of the aging-associated transcripts may predict protein levels well (r ≥ 0.5). In parallel, a comparison of the identified aging-regulated genes with public human transcriptomics data showed that only 35–45% of the identified genes show an age-dependent expression in corresponding human tissues. Finally, integrating both RNA-protein correlation and human conservation across data sources, we nominate 134 prioritized aging striated muscle signature genes that are predicted to correlate strongly with protein levels and that show age-dependent expression in humans. These prioritized signatures may hold promise to understanding heart and skeletal muscle physiology in human and mouse aging.

## Introduction

Skeletal and cardiac muscles are highly specialized tissues that are associated with distinct functional declines during aging. In aged organisms, there is a progressive loss of skeletal muscle mass, function, and regenerative capacity. Sarcopenia leads to frailty, diminishes the capacity for locomotion, limits the physiological role of muscles to regulate systemic glucose metabolism, and is a strong independent predictor of mortality in the elderly (Landi et al., 2013; Moore et al., 2014). In parallel, age-associated heart diseases are a leading cause of mortality and morbidity worldwide. Old age is associated with a decline in cardiac reserve, stress tolerance, metabolic and functional capacity, as well as the development of myocardial fibrosis that reduces the elasticity of the cardiac muscle (Lesnefsky et al., 2016; Triposkiadis et al., 2019). Hence, existing evidence strongly points to striated muscle tissues as being key to preserving organismal function and promoting healthspan. Understanding the molecular mechanisms of aging hearts and skeletal muscles is an important component in the quest to mitigate and prevent prevalent morbidities in an aging world.

Recent reports have surveyed the transcript level changes in the aging mouse skeletal muscle (Graber et al., 2021; Lin et al., 2018; Mikovic et al., 2018) or the heart (Bartling et al., 2019; Benayoun et al., 2019; Greenig et al., 2020). Separate studies have also determined the transcript (Timmons et al., 2019; Tumasian et al., 2021) and protein abundance changes (Murgia et al., 2017; Ubaida-Mohien et al., 2019a) in aging human skeletal muscles. Despite progress however, significant knowledge gaps persist. Continued investigations are needed to establish consistent aging signatures in the heart and the skeletal muscle across multiple models, and to contrast tissue-specific signatures across the two major groups of striated muscles. Moreover, important questions remain unanswered on whether and how much of the detected transcriptome changes might be translated to the protein level.

It is now established that transcript and protein levels correlate imperfectly across tissues and biological samples, where some transcripts may even be negatively correlated with the abundance of their protein counterpart (Franks et al., 2017; Jiang et al., 2020; Krug et al., 2020). Because proteins perform the overwhelming majority of biological processes, the results from transcriptomics data might be differentially relevant to biological processes based on how well they predict protein level. Recent large studies including GTEx (GTEx Consortium, 2020; Jiang et al., 2020), CPTAC (Krug et al., 2020; Mani et al., 2021), and GESTALT (Tumasian et al., 2021) have compared transcriptomics and proteomics data from matching tissues in large cohorts, and generally find moderate correlation between RNA and protein levels across samples. At present however, large-scale proteomics data of comparable depths remain far less accessible and common than transcriptomics data, hence there is intense interest in comparing across omics layers and identifying the transcriptomics signatures in aging and disease models that are translatable to the protein layer.

Here we performed total transcriptomic analysis in the heart and the skeletal muscle to assess global gene expression features of striated muscle aging. We apply a computational data analysis workflow that: (i) estimates the degree to which the transcriptome changes may predict changes at the protein level; and (ii) co-analyzes public human transcriptomics data to prioritized conserved signatures in human tissues. This approach may be useful for prioritizing transcriptomics signatures that are likely relevant to protein-level regulation.

## Experimental

### Animals and tissue extraction

All animal protocols were approved by the Institutional Animal Care and Use Committee at the University of Colorado School of Medicine. C57BL/6J mice were purchased from Jackson Laboratories (Bar Harbor, ME, USA) and housed in a temperature-controlled environment on a 12-h light/dark cycle and fed with normal diet and water ad libitum under National Institutes of Health (NIH) guidelines for the Care and Use of Laboratory Animals. Young adult mice (~4 months) and early aging mice (~20 months) mice (n=4, 2 male 2 female) were sacrificed, followed by measurement of body weight, heart weight and tibia length. The left cardiac ventricle and quadriceps femoris muscle were collected and stored at –80 °C.

### Total RNA sequencing

To extract RNA, the tissues were cut into ~1 mm^3^ cubes on ice. Cold TRIzol (Invitrogen) was added at 75 μL per mg tissue and tissues were homogenized on a bead mill homogenizer with 2.8 mm ceramic beads at 10 second duration, speed 5, 5 repeats. The samples were centrifuged at 16,000 g for 15 minutes at 4 °C and the supernatant was transferred to RNase-free tubes. RNA extraction was performed using the Direct-zol RNA Miniprep Plus kit (ZYMO) following manufacturer’s instructions. Total RNA sequencing was performed on the tissues (~80M reads/ 20 Gbases, 151 nt PE, Zymo Ribodepleted library) using Illumina short-read sequencing on a NovaSeq 6000 platform. The data were mapped to the mouse genome GRCm38.p6 using STAR v.2.7.6a (Dobin and Gingeras, 2016). The mapped transcripts were assembled using Stringtie v.2.1.1 (Kovaka et al., 2019) against Gencode vM25 gff3 annotations. All sequencing data are available on GEO at GSE175854.

### Liquid chromatography and mass spectrometry

To extract proteins, tissue pieces were lysed in RIPA buffer and Halt protease/phosphatase inhibitor (Thermo) with a hand-held homogenizer, followed by sonication and centrifugation at 16,000 g for 15 minutes at 4 °C. Protein quantity was measured using BCA assay (Thermo), after which 100 μg of proteins were digested using a filter-assisted protocol as described (Manza et al., 2005). Digested peptides were desalted using C18 spin columns (Pierce). Label-free bottom-up mass spectrometry was performed using data dependent acquisition on an Orbitrap Q-Exactive HF connected to a Easy-nLC 1200 nano-UPLC system using typical settings as described (Lau et al., 2019). Mass spectrometry raw data was converted to mzML using ThermoRawFileParser v.1.2 (Hulstaert et al., 2020), and searched against UniProt SwissProt (The UniProt Consortium, 2018) reviewed *Mus musculus* database (retrieved 04/27/2021) with appended contaminant sequences using MSFragger v.3.2 (Kong et al., 2017), followed by post-processing using PeptideProphet and ProteinProphet in the Philosopher suite v.3.4.13 (da Veiga Leprevost et al., 2020) and label-free quantification with match-between-runs using IonQuant v.1.5.5 (Yu et al., 2020).

### Retrieval and analysis of public expression data sets

Human gene expression profiles are from GTEx v8 release (GTEx Consortium, 2020) data retrieved from the GTEx portal and normalized using variance stabilizing transformation (Love et al., 2014), then batch corrected with the aid of ComBat (Leek et al., 2012). CPTAC RNA-seq and mass spectrometry datasets for breast (Krug et al., 2020), ovarian (Hu et al., 2020b; Zhang et al., 2016), colorectal (Vasaikar et al., 2019; Zhang et al., 2014), lung adenocarcinoma (Gillette et al., 2020), and endometrial (Dou et al., 2020) cancer discovery studies were retrieved in accordance with the CPTAC data use and embargo policies using the cptac v.0.9.1 package in Python 3.9. Statistical learning was performed using scikit-learn 0.24.2 (Lindgren et al., 2021). Transcriptomics data were standardized, after which data were split 80/20 into train and test sets. Prediction was performed using an elastic net for the sake of consistency with the CPTAC DREAM baseline model (Yang et al., 2020). Correlation coefficients, R^2^, and normalized root mean square error metrics were reported.

### Additional statistics and data analysis

Data analysis was performed in R v.4.0.5 and Bioconductor v.3.12 (Huber et al., 2015) on an x86_64-apple-darwin17.0 (64-bit) platform. Statistical tests for sequencing reads were performed using DESeq2 v.1.30.1 (Love et al., 2014) using age and sex as factors after filtering low-read genes to retain 17,003 genes in the heart and 16,231 genes in the muscle. Statistics for label-free proteomics data were performed using limma v.3.46.0 (Ritchie et al., 2015) with age and sex as factor. Cell type proportions were estimated with the aid of Seurat v.4.0.1 (Butler et al., 2018), the weighted nonnegative least square regression method in MuSiC v.0.1.1 (Wang et al., 2019), and Tabula Muris FACS mouse heart and muscle single-cell RNA sequencing data (Tabula Muris Consortium et al., 2018). Functional enrichment was performed with the aid of fgsea v.1.16.0 (Sergushichev, 2016) and MSigDB v.7.3 annotations (Liberzon et al., 2011) or ReactomePA v.1.9.4 (Yu and He, 2016) against Reactome annotations (Fabregat et al., 2018) loaded in the package. Comparison of correlation coefficients was performed using cocor v.1.1-3 (Diedenhofen and Musch, 2015).

## Results & Discussion

### Global features and pathways in aging heart and muscle

We first acquired total RNA sequencing data on the total transcriptome changes in the heart and the skeletal muscle between young adult vs. early aging mice (4 vs. 20 months) (n=4 each) (**Supplementary Table S1**). The transcriptome profiles in each organ is distinguishable by sex, and shows more separation by ages in female than in male animals (**Fig. 1A**). On a global level, we considered the cellular pathways that are involved with age-associated gene expression changes using the fast gene set enrichment analysis (FGSEA) algorithm. In both tissues, aging is associated with a positive enrichment of genes involved in innate immune system and neutrophil degranulation (FGSEA P.adj 1.4e–3 heart; 1.0e–2 muscle) as well as GPCR ligand binding terms (P.adj 8.5e–2 heart; 3.1e–3 muscle); and a negative enrichment of genes involved in the respiratory chain (P.adj 3.8e–3 heart; 6.7e–3 muscle), citric acid cycle (P.adj 4.2e–3 heart; 8.1e–3 muscle), and translation (P.adj 6.9e–2 heart; 1.1e–2 muscle) (**Figure 1B**). Comparable processes were enriched using WikiPathways and KEGG terms as annotations (**Supplementary Figure S1**). A major discrepancy between the two types of muscle involves genes functioning in extracellular matrix organization and collagen degradation, which are up-regulated in aging hearts but down-regulated in aging tissues.

**Figure 1:**
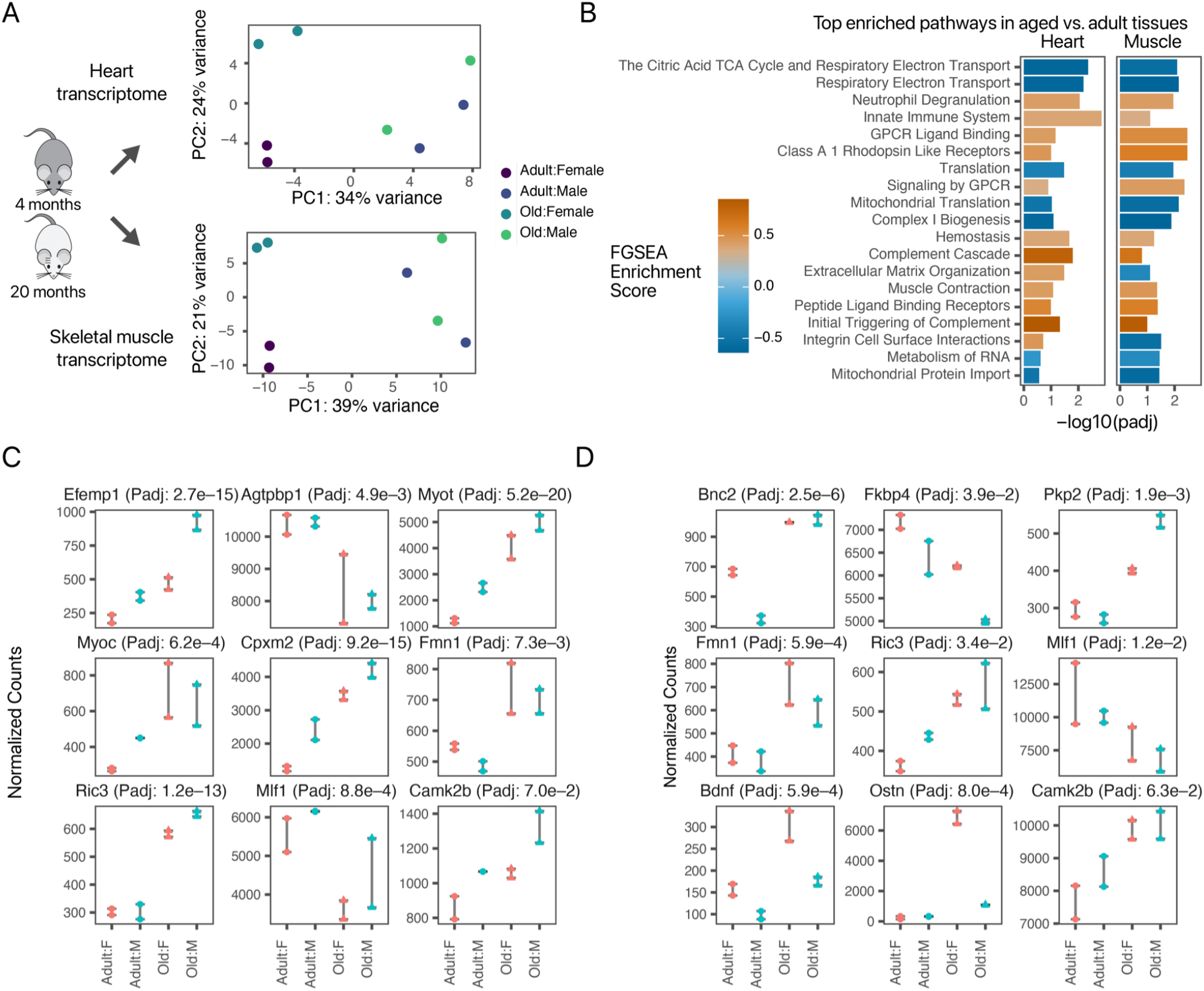
Transcriptome changes in young adult mice vs. early aging mice. **A.** Principal component analysis of normalized gene counts in the left ventricle (top) and quadriceps femoris (bottom) show separation by sex and age. **B.** Enriched pathways in aged vs. adult gene expression in two tissues. **C-D.** Normalized read counts showing sex and age expression among selected differentially expressed aging genes in (**C**) the heart and (**D**) the skeletal muscle (FDR 10%).

Closer inspection of the fold-changes of genes making up the leading edges of FGSEA-enriched terms show a concomitant down-regulation of genes in electron transport and translation pathways and up-regulation of innate immune system genes (**Supplementary Figure S2A–C**). Although muscle contraction genes are implicated in both tissues during aging, we find that the up-regulated genes differ, with *Sln, Myl7*, and *Myl4* most prominently induced in the heart as opposed to *Myh6* and *Myl2* in the skeletal muscle (**Supplementary Figure S2D**), whereas we also observed individual genes in extracellular matrix organization changed in opposite directions as the pathway enrichment results suggest (**Supplementary Figure S2E**).

Taken together, the pathway-level analysis suggests that during normal heart and skeletal muscle aging, gene expression changes are consistent with a rerouting of gene expression from mitochondrial metabolism and protein synthesis usage toward inflammatory and matricellular functional components, although the changes in extracellular matrix appeared to diverge between the two striated muscle. Our results also add to a chorus of recent findings that implicate the innate immune system in muscle aging (Graber et al., 2021; Lin et al., 2018; Ubaida-Mohien et al., 2019b) and other tissues (Benayoun et al., 2019).

### Transcriptomic signatures in striated muscle aging

We next considered the signatures implicated in aging at an individual gene level. We found 358 differentially expressed coding genes in old vs. adult hearts and 276 in skeletal muscles of identical animals at 10% FDR (**Supplementary Data S1–S2**). Among the induced genes, we found genes that are previously associated with age-associated diseases as well as genes not previously associated with aging tissues. Among the genes of interest, nuclear receptor subfamily 4 group A member 1 (*Nr4a1*) and mothers against decapentaplegic homolog 3 (*Smad3*), were both significantly decreased in aging skeletal muscle (*Nr4a1* logFC –0.73, P.adj 3.8e–4; *Smad3* logFC –0.44, P.adj 2.1e–2). A recent large-scale meta-analysis of 739 human skeletal muscle transcriptomes from endurance or resistance exercise interventions pinpointed the human *SMAD3* and *NR4A1* genes as a central hub of acute response to exercise in the skeletal muscle (Amar et al., 2021). Both *SMAD3* and *NR4A1* are acutely up-regulated after exercise, contradirectional to the observed age-associated repression here. *Smad3* interacts with *Stat3*, which in skeletal muscle may function to regulate muscle mass (Chao et al., 2012), whereas *Nr4a1* may function to regulate mitochondrial biogenesis (Chao et al., 2012), hinting at potential connections of these genes to the benefits of exercise in delaying age associated muscle and mitochondrial loss.

Results on an individual-gene level were less conserved with other studies that compared young and aging mice than pathway changes. For instance, Lin et al. (Lin et al., 2018) and our data both found strong changes in immune regulation, but Lin et al. reported a strong decrease in *Fkbp5* in aging skeletal muscle which was not recapitulated in this study. Instead, we found a significant decrease in *Fkbp4* (**Figure 1D**) in aging skeletal muscle. Similarly, although both work noted an upregulation of muscle contraction genes in aging, the specific genes overlap only partially, with Lin et al. reported upregulation of *Myh7*, *Myh3*, *Tnnt1*, among others, and the present data are represented by *Myo5a, Tpm2*, and *Tnnt2* (**Supplementary Data S2**). Lastly, among the top 20 positively age-correlated and top 20 negatively age-correlated human genes in the NIA GESTALT study (Tumasian et al., 2021), we observed evidence for 5 being recapitulated here (up-regulated: *Skap2, Cfap61, Kcnq5;* down-regulated: *Myl1, Casq1* at 15% FDR). These across-study differences can plausibly arise from the use of arbitrary significance cutoffs as well as a combination of differences in study design, organism models, technical variations, and stochastic gene regulations. We hypothesize that both individual-gene level and pathway level changes contain complementary information into the molecular mechanisms of aging, which should be taken into account when multiple studies are compared in meta-analyses.

Interestingly, we identified a constellation of genes coding for secreted proteins, which was not the focus of prior reports in aging mice. For instance, in the heart, inactive carboxypeptidase-like protein X2 (*Cpxm2*) encodes a secreted protein that is induced in aging hearts (logFC 1.1, DESeq2 P.adj 9.1e–15). Likewise induced is EGF-containing fibulin-like extracellular matrix protein 1/fibullin 3 (*Efemp1*) (logFC 1.25, P.adj 2.7e–15), which encodes an extracellular matrix protein that may be cleaved into a secreted peptide and that binds with EGF receptor (**Figure 1C**). Very recently, the NIA GESTALT study has also found human EFEMP1 to be positively correlated with age in the skeletal muscle (Tumasian et al., 2021). In the normal aging muscle, osteocrin (*Ostn*) encodes a secreted hormone musclin that acts as an exercise induced myokine (Subbotina et al., 2015) but also functions in the heart where it may protect against apoptosis and inflammation (Hu et al., 2020a). In the analyzed animals, *Ostn* is induced in normal aging animals but most prominently in females (**Figure 1D**) (logFC 3.3, P.adj 8.0e–4). *Bdnf* encodes a myokine that is induced by exercise and regulates energy metabolism at least in female mice (Yang et al., 2019); we found dimorphic expression with higher expression in female and which is further induced in aged tissues (logFC 0.9, P.adj 5.9e–4). Other secreted factor encoding genes changed in aging include *Gdf11* (logFC 1.76, P.adj 9.5e–2) in skeletal muscle as well as *Vegfd* (logFC 0.85, P.adj 5.8e–2), *Frzb* (logFC 0.69 P.adj 9.3e–2), *Sfrp1* (logFC 0.62, P.adj 6.6e–3), *Fstl4* (logFC 0.57, P.adj 3.6e–3), and *Fgf13* (logFC –0.27, P.adj 4.8e–2) in the heart (**Supplementary Data S1–2**).

We identified 20 common coding genes that are differentially expressed in aging in both tissues at 10% FDR (**Fig. 2A**). The shared genes show strong positive correlations (Spearman’s correlation coefficient r 0.65, P 0.0032) with the exception of one outlier (predicted gene *Gm50364*), which is repressed in the muscle and induced in the normal aging heart (**Fig. 2B**). Among the common genes, formin-1 (*Fmn1*) codes for a myofibril differentiation factor that plays a role in the formation of adherens junction and is increased in aged muscles and hearts. Myeloid leukemia factor 1 (*Mlf1*) codes for a protein that may serve as a negative regulator of cell cycle exit and is suppressed in both aging hearts and muscles.

**Figure 2:**
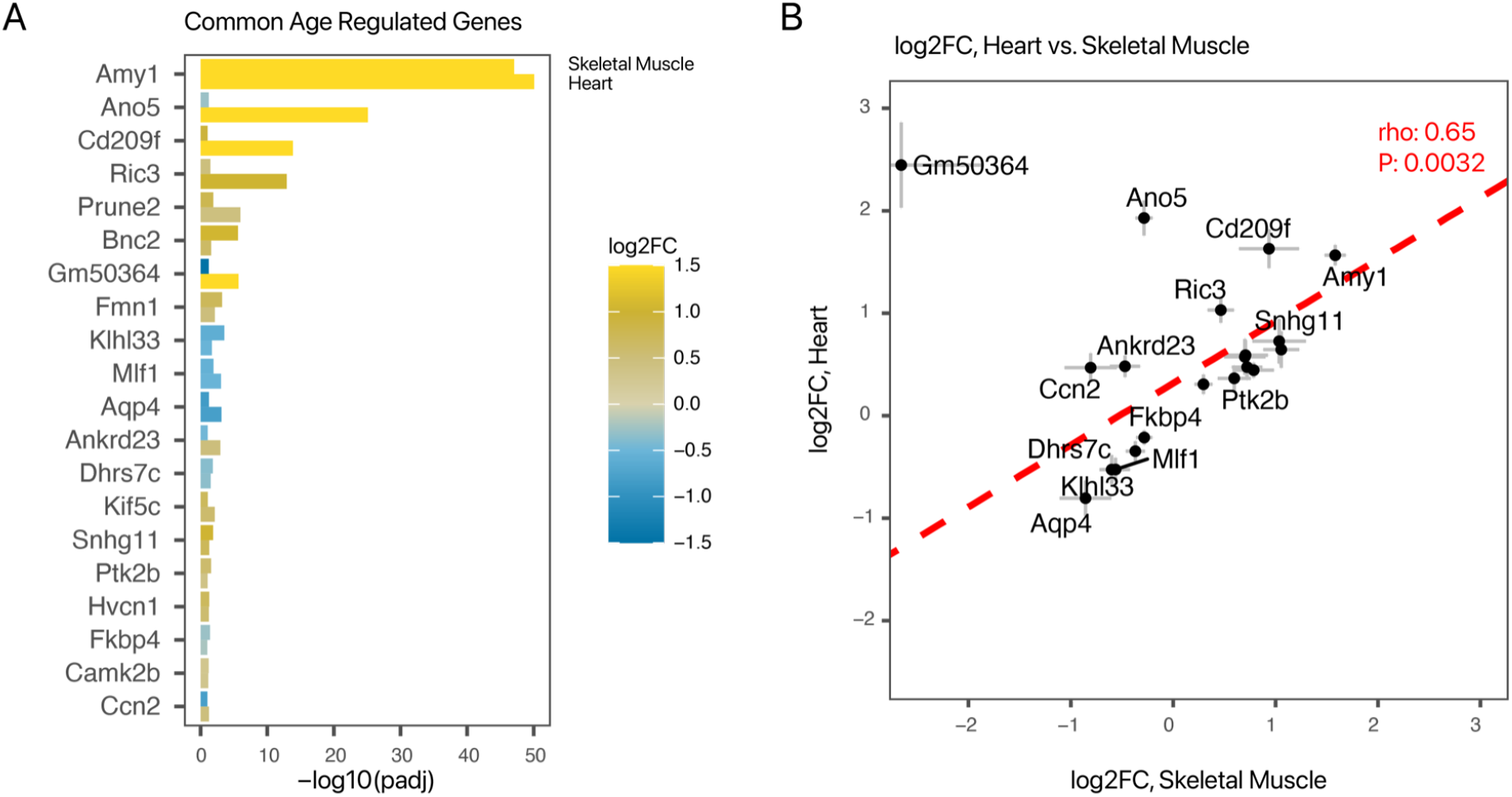
Shared aging signatures in aged cardiac and skeletal muscle. **A.** Bar chart showing the adjusted P values (x-axis) and log2 fold-changes (log2FC; fill color) of the 20 aging associated genes identified in both tissues. **B.** Scatter plot showing a comparison of fold changes (aged vs. young adult) and a robust positive correlation (Spearman’s correlation coefficient ρ 0.65, P 0.0032) between the two tissues. X-axis: log2 fold-change in skeletal muscle; y-axis: log2 foldchange in the heart. Error bars: standard error of log2 fold-change; line: best-fit linear curve.

To assess whether the gene expression changes were due to changes in cell type compositions, we decomposed the bulk RNA-seq read count matrices into individual cell types derived from single-cell sequencing data using weighted non-negative least square methods (**Supplementary Figure S3**). We found no evidence of substantial changes of overall cell type proportion, suggesting the observed transcriptomic changes are unlikely due to wholesale changes in cell population in young adult vs. aged hearts and muscles.

### RNA-protein correlation and predictability of protein-level changes

To estimate whether the effect of aging transcriptome changes is potentially translated to proteomic changes, we first applied a statistical learning method against one of the largest proteogenomics data sets in existence to train a model to predict protein levels using their cognate transcript levels as proxy (see **Methods**). The correlation coefficients, R^2^ values, and normalized root mean square errors (NRMSE) values between the transcript-predicted protein level and empirical mass spectrometry-measured protein levels across subjects are taken as the protein predictability of a transcript for each gene. In total, we estimated the protein predictability of 11,896 transcripts (**Supplementary Data S3**). We found a large range of predictability where the predicted-actual Pearson’s correlation coefficients ranged from –0.822 to 0.999 (interquartile range 0.18–0.54) (**Figure 3A-B**). A small portion of transcripts (4.5%) had negative correlation with protein levels. The median r of all genes is 0.374 which is comparable to previously reported RNA-protein correlation in comparable data sets (Eicher et al., 2019; Li et al., 2019; Yang et al., 2020) and in human tissues (Jiang et al., 2020).

**Figure 3:**
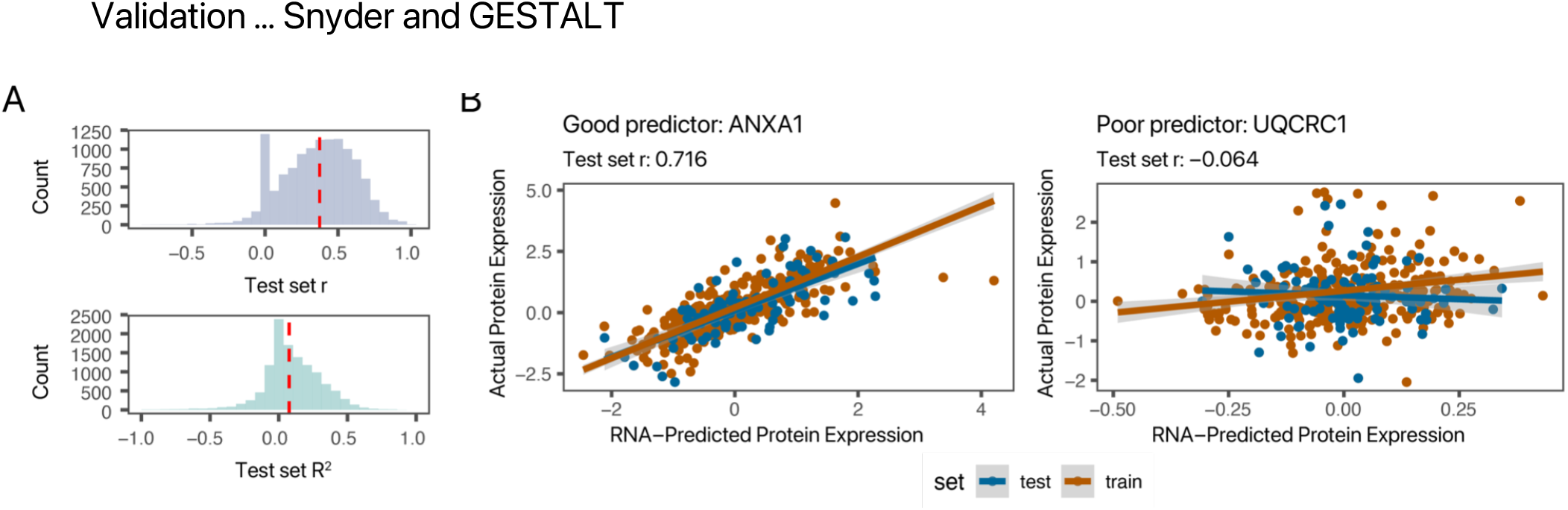
Predictability of protein-level from across-sample transcript variance. To estimate whether the quantified transcript changes might translate to the proteome, we considered the predictability of protein levels from their proxy transcripts on a gene-wise basis in large proteogenomics datasets. **A.** An elastic net is applied to 717 samples with matching transcriptomics and mass spectrometry data in the CPTAC collection. The average correlation (top) and R^2^ values (bottom) between predicted and actual protein levels across samples in each of 10,693 genes are shown. **B.** Examples of an aging signature whose protein abundance across samples is well predicted by its proxy transcript (*Anxa1*) in matching samples and one that is poorly predicted (*Uqcrc1*). Each data point is one CPTAC sample. Brown: train set; blue: test set.

We found that poor predictors (r ≤ 0.3) are enriched in pathways involving major multiprotein complexes, including Reactome Translation (P.adj 1.9e–62), mRNA Splicing (P.adj 4.4e–24), and Respiratory electron transport (P.adj 2.8e–15) terms. Good predictors (r ≥ 0.7) are enriched in Reactome Biological oxidations (P.adj 1.9e–6), Extracellular matrix organization (P.adj 2.1e–4), and Metabolism of lipids (P.adj 7.8e–3) terms (**Supplementary Data S4**). This agrees with emerging themes from cancer and normal tissue studies. For instance, a large-scale GTEx survey of 32 normal human tissues has found that secreted proteins and proteins in multi-protein complexes are generally poorly predictable from transcripts (Jiang et al., 2020), presumably because of additional post-translational constraints on their steady state levels. This corroborates that different proteins exhibit a wide range of predictability by proxy transcripts, and hence different transcripts have different intrinsic value in reflecting actual protein abundance states across tissues, and moreover, predictions of RNA-protein agreement might be transferable across to different samples and based on basic biophysical constraints. For example, long-half-life housekeeping proteins are usually more predictable from transcripts whereas the abundance of multi-protein complex members are buffered by complex stoichiometry and assembly.

To further verify the potential impact of the transcriptome signatures on the proteome, we performed in-house exploratory proteomics analyses of identical tissues from identical animals using label-free quantitative tandem mass spectrometry (**Supplementary Data S5**). In total, we acquired the MS1 label-free quantity of 1,254 distinct proteins in the heart and the skeletal muscle identified with ProteinProphet protein probability ≥ 0.95. We identified 58 and 76 proteins with nominal limma P ≤ 0.05 and |logFC| ≥ 0.5 in the heart and the skeletal muscle, respectively, although only 6 proteins reached adjusted P ≤ 0.1, presumably due to the limited depth and breadth of the proteomics profile performed here. Nevertheless, pathway analysis using FGSEA against MSigDB revealed an enrichment of similar annotation terms to the transcriptomics data, including Reactome Extracellular matrix organisation (FGSEA permutation P: 3.5e–4) and Innate immune system (P: 0.053) terms, suggesting the mass spectrometry experiment was able to capture a representative footprint of the aging proteomes in these tissues.

Not unexpectedly, we found there is a robust correlation between RNA and protein relative abundance *across* genes *within* a tissue (Pearson’s r: 0.51 heart; 0.46 muscle) (**Figure 4A**). There was a modest decrease in correlation of RNA and protein levels in aged samples as previously reported, but in our data this difference was not significant (Fisher’s z P: 0.49 heart 0.46 muscle). The correlation between RNA and proteins weakens significantly when correlations across samples are considered (Pearson’s r: 0.18 heart; 0.14 muscle) (**Figure 4B**). This reflects the distinction between RNA-protein correlation across genes vs. across samples, and corroborates mounting evidence that show although abundant proteins tend to have abundant transcripts, transcript changes are imperfectly correlated to proteins due to post-transcriptional activation (Franks et al., 2017). Nevertheless, when only age-differentially expressed transcripts (10% FDR) were compared, we observed a general concordance in the directionality of protein changes (**Figure 4C**). Notably, RNA-protein correlation is higher among transcripts with nominal changes (DESeq2 P ≤ 0.1) that had higher estimated protein prediction (1 ≥ r ≥ 0.5) than those with lower prediction (0 ≤ r ≤ 0.5) (correlation 0.42 vs. 0.11; cocor P 1.5e–3) (**Supplementary Figure S4**). The analysis therefore corroborates the transferability of the learned model and offers supportive evidence for the adaption of predicted RNA-protein correlations as one optional method to help prioritize discovered transcript signatures.

**Figure 4:**
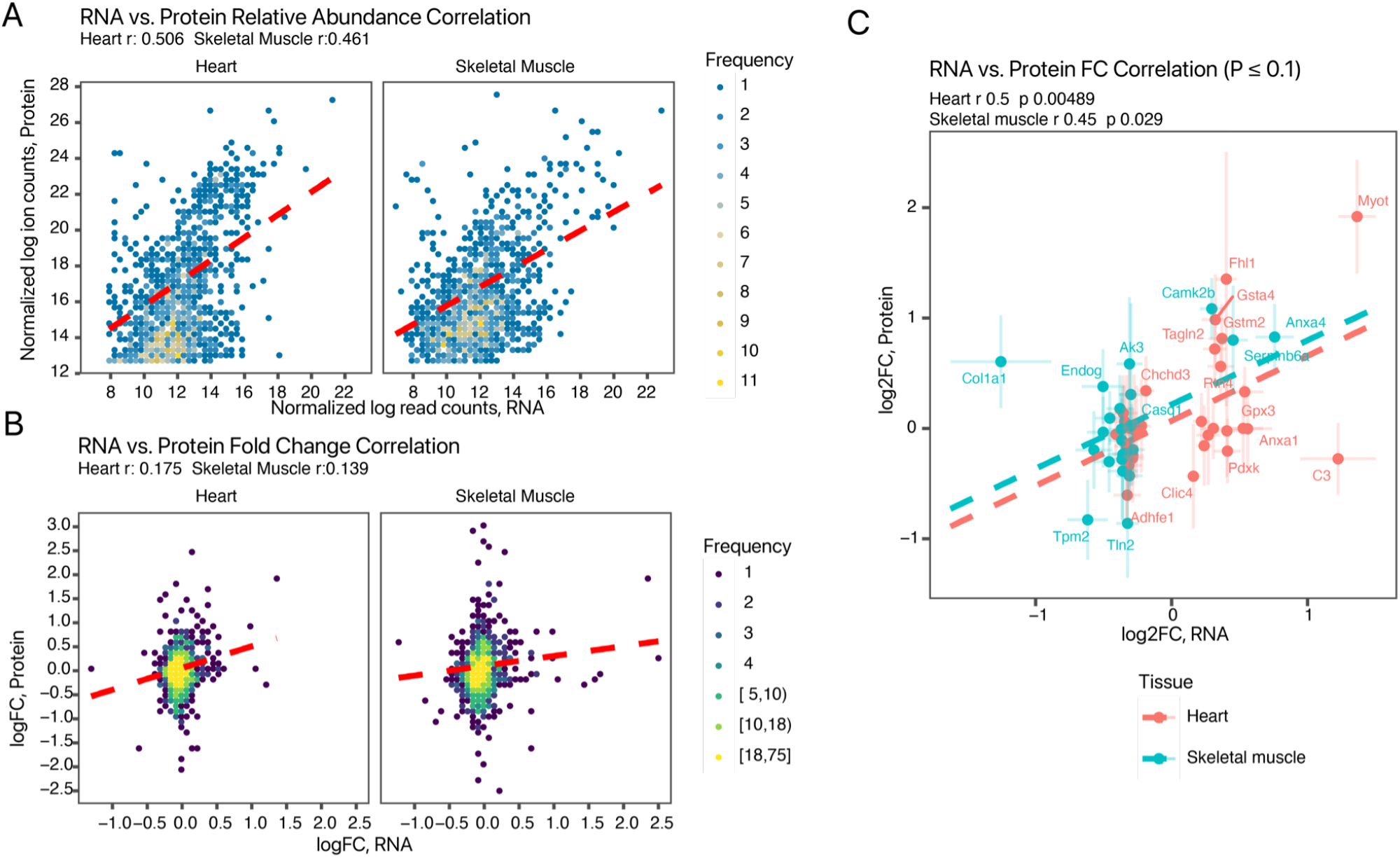
Correlation between RNA and protein levels from identical tissues. **A.** Scatter plot showing the within-sample across-gene comparisons in the heart (left) and the skeletal muscle (right) for commonly quantified RNA and their proteins. Fill color: data frequency within bin. **B.** Scatter plot showing across-sample comparisons between RNA and proteins in the aging vs. young adult heart (left) and skeletal muscle (right). **C.** Log fold-change comparison at the RNA (x-axis) and protein (y-axis) level among commonly quantified proteins and transcripts with significant age-associated transcript level differences. Line: best-fit linear curves for the heart (red) and the skeletal muscle (blue). Error bars: standard errors of logFC.

### Conservation of identified striated muscle aging signatures in human

Because specific genes and pathways may underlie aging processes in different organisms, we next estimated the extent to which the identified mouse aging transcriptome features are translatable to humans. To do so, we analyzed the age vs. expression relationship of GTEx v8 human transcriptomics data. In total, we retrieved 17,382 RNA sequencing samples including 432 heart left ventricle, 429 atrial appendage, and 803 skeletal muscle transcriptomes, and performed stepwise normalization for technical, tissue, and donor batches (**Supplementary Figure S5A–E**). We then compared whether the normalized gene expression of each gene signature is correlated with donor age groups in different tissues in humans. We found overall there are complex trends between identified age-associated signatures with human gene expression-age group relationship. Only 35% (107/305) and 44% (97/217) of the analyzed aging genes in the mouse heart and the skeletal muscle, respectively, had significant age-expression relationship in GTEx v8 (ANOVA P ≤ 0.01), suggesting not every identified signature is potentially conserved across species (**Supplementary Figure S6**). On a global level, up-regulated genes in normal aging mouse hearts are significantly more likely to be positively correlated in expression with donor age groups in human heart left ventricle (Wilcoxon P: 8.0e–5) and atrial appendage (P: 2.8e–2) samples but not in the other compared human tissues including the kidney cortex (P: 0.33) or liver (P: 0.27). Moreover, this correlation is not existent when only sexually dimorphic (sex-differentially expressed at 10% FDR) genes in the mouse are compared in human hearts (Wilcoxon P: 0.87 for heart - atrial appendage and 0.95 for heart - left ventricle). This global relationship is considerably subdued for genes that are differentially regulated in aging muscle, which may suggest that the aging signatures in this tissue are more specific to species or otherwise show non-linear change over the lifespan (**Supplementary Figure S5**).

Incidentally, the age-expression relationships of signature genes are often not preserved in other tissues despite the gene being expressed at appreciable levels. For example, *Efemp1* is induced in aging in the mouse data here, and is positively correlated with age group in GTEx v8 human heart and skeletal muscle tissues but not in the kidney or the liver, despite the human *EFEMP1* gene being expressed at a similar baseline level in those tissues (**Figure 5A**). To corroborate this incidental observation, we acquired and analyzed RNA sequencing data from the kidney of identical animals (**Supplementary Data S6**). The data corroborated that there are no significant changes in *Efemp1* (logFC 0.20, P.adj 0.28). Likewise, *Fkpb4* is not significantly changed in the kidneys of aging humans in GTEx v8 or the RNA-seq data of identical animals (logFC 0.03, P.adj 0.92) (**Figure 5B**); *Sod3* is correlated with age in human hearts and muscles but not kidney, whereas we also found no significant changes in mouse kidney in our RNA-seq data (logFC 0.00, P.adj 1.00) (**Figure 5C**). Taken together, the results suggest that normal aging signatures exhibit tissue specificity, both across skeletal and cardiac muscles as well as between striated muscles and other organs, as well as potential species specificity, and a selection strategy might be employed to prioritize aging signatures that show expression trends in humans.

**Figure 5:**
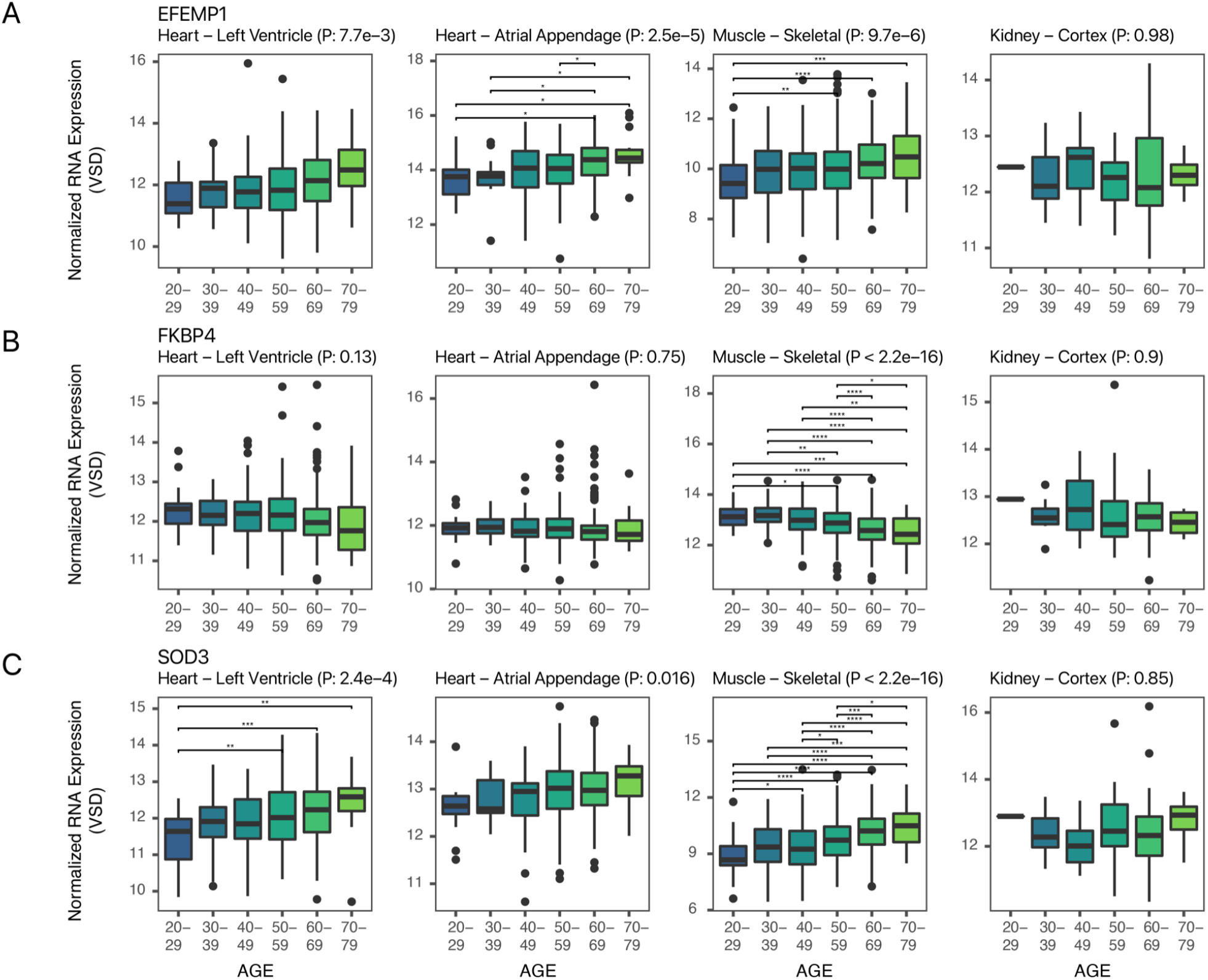
Conserved age-expression profiles of selected signatures in humans. Box plots showing GTEx v8 human normalized RNA expression levels across age groups in decadal brackets in four GTEx v8 human tissues (left to right) heart left ventricle, heart atrial appendage, skeletal muscle, and kidney cortex for **A.***Efemp1*,**B.** *Fkbp4*, and **C.** *Sod3*. P values: ANOVA. Asterisks within plots denote Tukey’s post-hoc for individual group comparison. *: Tukey P < 0.05; **: P ≤ 0.01; ***: P ≤ 0.001; ****: P ≤ 0.0001.

Integrating information from both RNA-protein correlation and human conservation, we ranked the identified aging signatures based on how well the transcript-predicted across-sample protein values reflect empirical protein levels (r ≥ 0.5), and moreover selected signatures that show a significant correlation with age group in human GTEx v8 tissues (ANOVA P ≤ 0.01). This combined analysis led to 134 of the prioritized age-associated signatures that are potentially more likely to have bonafide relevance in the biology of aging tissues at the protein levels (**Figure 6**).

**Figure 6:**
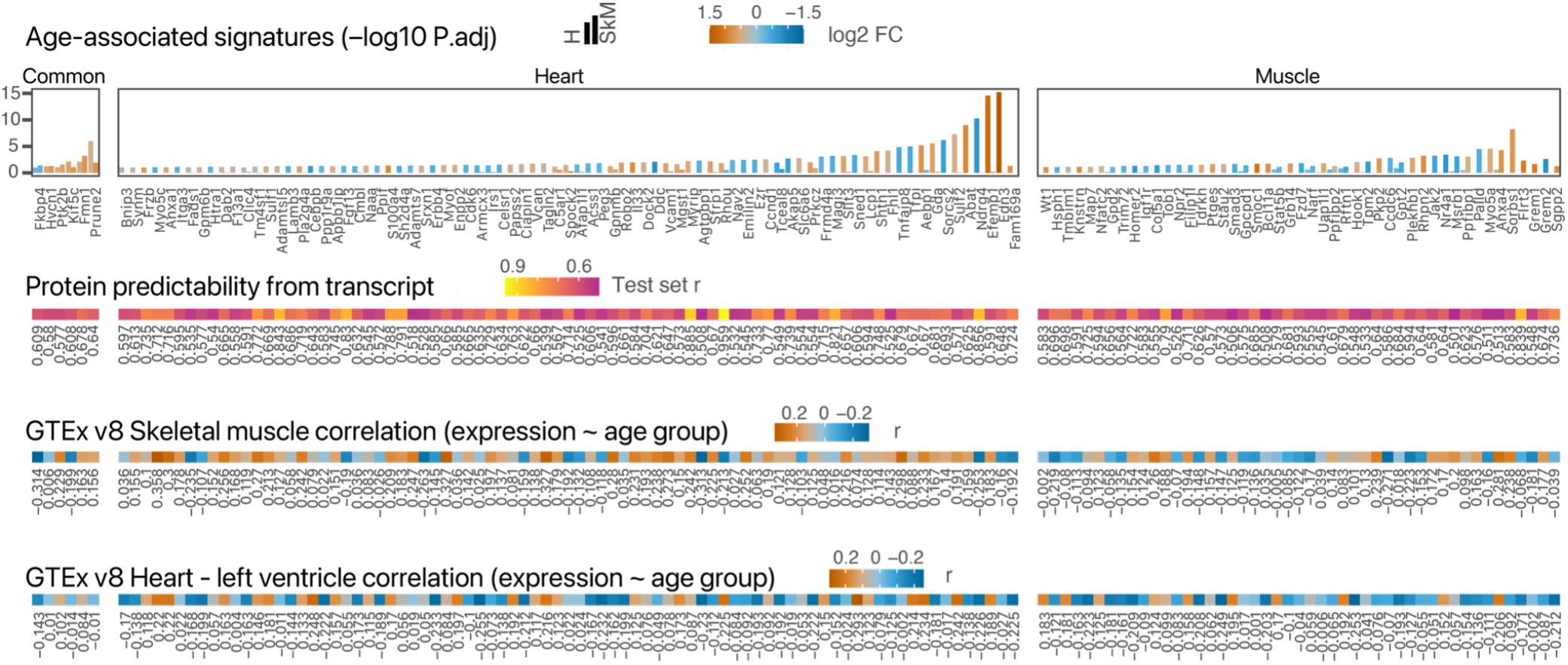
Prioritized age-associated signatures. List of 134 prioritized aging signatures in the heart, skeletal muscle, and those common to both tissues. Top: paired bars represent –log10 P.adj in the heart and the skeletal muscles, respectively. Row 2–4: the prioritized signatures had CPTAC RNA-protein correlation r ≥ 0.5 and ANOVA P ≤ 0.01 in GTEx v8 transcript expression against age groups in GTEx v8 heart left ventricle or skeletal muscle transcriptomes.

### Non-coding genes in striated muscle aging

Lastly, although the focus of this study is to prioritize protein correlation of coding gene signatures, not all transcripts function only through their translation products, and non-coding RNAs play important roles in virtually all aspects of biology. As we acquired total ribosomal-depleted RNA abundance data, we also explored the changes of non-coding RNAs in aging hearts and muscles including non-poly-A+ transcripts. From the data, we found 19 noncoding genes to be differentially expressed with age in the heart and 36 in the muscle at 10% FDR, with 3 overlapping. Grouping into gene category annotations suggests that the differentially expressed non-coding genes included sense, anti-sense, and intergenic long non-coding RNAs (lncRNAs), as well as small nucleolar RNAs and processed pseudogenes (**Figure 7A**). The non-coding RNAs were manually inspected for read mapping and strand specificity. We recapitulated changes in two maternally imprinted lncRNAs *Meg3* and *Riat*, which were previously found using qPCR array to be decreased in skeletal muscle over the lifespan (Mikovic et al., 2018)), although *Meg3* was also found to be increased in senescent endothelial cells (Boon et al., 2016). We also identified additional age-regulated lncRNAs. In the muscle, *Plet1os* is also decreased (logFC –1.41, P.adj 8.8e–10) (**Figure 7B**), whereas *Foxo6os* is decreased in aging (logFC –0.63, P.adj 0.002), and was previously found to be depressed in insulin resistant muscle (**Figure 7C**). In the heart, the lncRNA *Mhrt* is located on the opposite strand of mouse *Myh7* and has been associated with the regulation of *Myh6/Myh7* ratios as well as protection against pathological cardiac remodeling (Han et al., 2014); in the data, we found *Mhrt* to be drastically reduced (logFC – 0.44, P.adj 2.7e–7) in aged hearts. Among the age-regulated lncRNAs, five (*Neat1, Plet1os, Foxo6os, Peg13, Mhrt*) were previously found to have potential translatability in smProt (Hao et al., 2018) or engaged in ribosomes in the mouse heart (van Heesch et al., 2019), suggesting a possibility that they may be translated. A growing number of microproteins are known to be translated in striated muscles. Future work combining transcriptomics and proteomics approaches might determine whether they are differentially regulated in aging or participate in associated pathophysiological processes.

**Figure 7.**
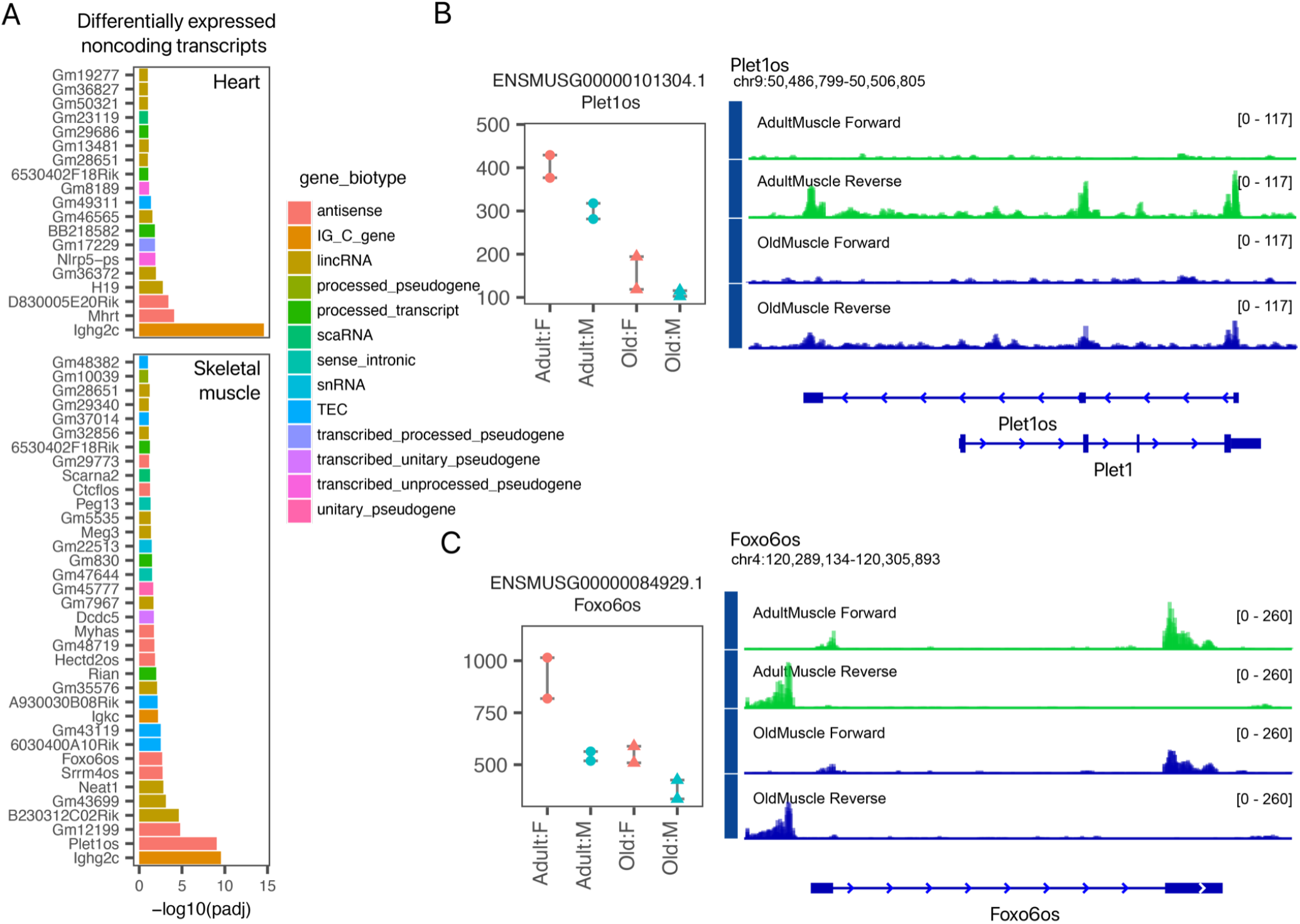
Non-coding RNA signatures in aging striated muscles. **A.** Bar charts of differentially expressed annotated non-coding RNAs in the heart (top) and the skeletal muscle (bottom). Colors denote Gencode vM25 annotation gene biotype. **B-C**. Examples and genome tracks of two long non-coding RNAs *Plet1os* (**B**) and *Foxo6os* (**C**) that are differentially expressed in aging skeletal muscle.

## Conclusion

This study examined the aging transcriptome profiles in skeletal and cardiac muscles. In total, we identified over 600 differentially expressed transcripts in the heart and the skeletal muscles of 20 months vs. 4 months aged mice. Our results add to mounting evidence that point to extracellular matrix, mitochondrial, and innate immunity processes as distinguishing factors in aging tissues. Notably, our results also point to a number of age-associated gene encoding for secreted signals, suggesting age-associated myokines and cardiokines present a promising avenue for further understanding the molecular mechanisms of heart and muscle aging. A number of identified signatures are specific to striated muscles aging while unchanged over age in other tissues, and hence may be particular to muscle aging processes.

We devised a computational workflow that transfers the predicted correlation between transcript and protein levels trained from a large data set as a means to prioritize potential age-associated signatures in a current small-scale data. About 48% of identified aging signatures are predicted to correlate well with the abundance of their protein counterparts. The performance of this transfer learning approach is expected to benefit from the continued accrual of additional data in closely related tissues and species. Further comparison to public human transcriptomes data showed that only 35–45% of the aging signatures show age-dependent expression in corresponding human tissues, prompting us to prioritize a subset of signatures that may be more likely conserved in humans. We suggest that this computational data analysis approach may be applied to future transcriptomics studies in aging and disease to help prioritize potentially biologically relevant and human translatable signatures.

Finally, there are several limitations pertaining to the experimental results here. For instance, the aging animals (20 months) used here are comparatively younger than the C57BL6/J mice in some other studies (22–28 months) (Bartling et al., 2019; Graber et al., 2021; Greenig et al., 2020; Mikovic et al., 2018), which might decrease the sensitivity of age-associated signatures and omit signals that only appear in very elderly animals (Graber et al., 2021). Second, only a label-free proteomics experiment of limited depth was performed as our major goal was to compare transcriptomics and proteomics fold change. Future work using deep quantitative proteomics comparison with common stable isotope labeling mass spectrometry techniques may help reveal specific proteomics features of aging that are not apparent at the transcript level, or identify non-canonical translation products using proteogenomics methods.

## Acknowledgments

This work was supported in part by NIH awards F32-HL149191 to YH; the UC Denver MARC U-STAR training program T34-GM096958 (NLK); R21-HL150456, R00-HL127302, and R01-HL141278 to MPL; and R00-HL144829 to EL. Data utilized in this publication include those generated by the National Cancer Institute Clinical Proteomic Tumor Analysis Consortium (CPTAC).

## Supplementary Figures and Tables

**Supplementary Figure S1.**
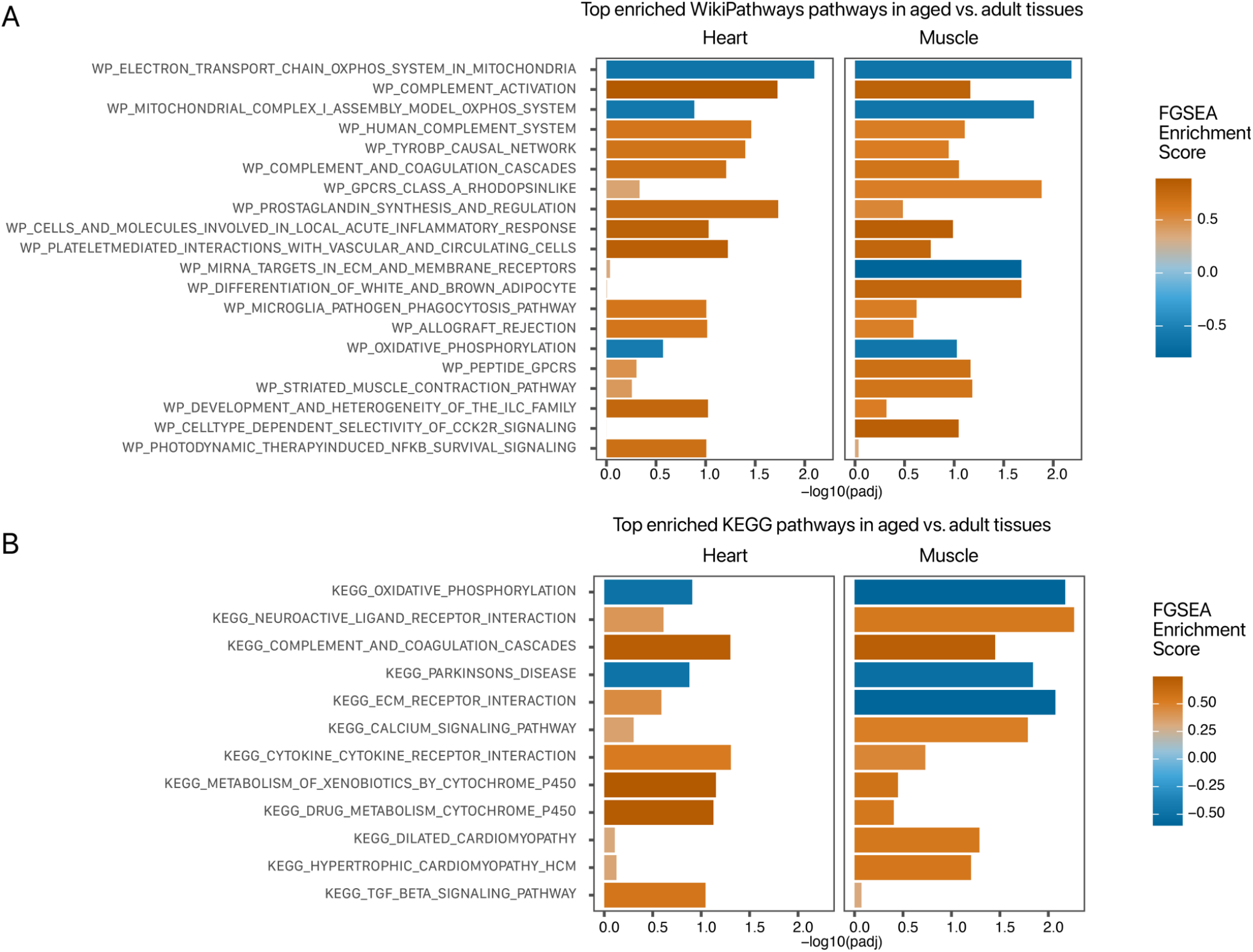
Additional aging associated pathways. **A–B.** Significantly enriched pathways in aging heart and skeletal muscles against **A.** WikiPathways and **B**. KEGG annotations. Pathway names are printed as given in MSigDB files. X-axis: –log10 FGSEA P.adj values; fill color: enrichment score.

**Supplementary Figure S2.**
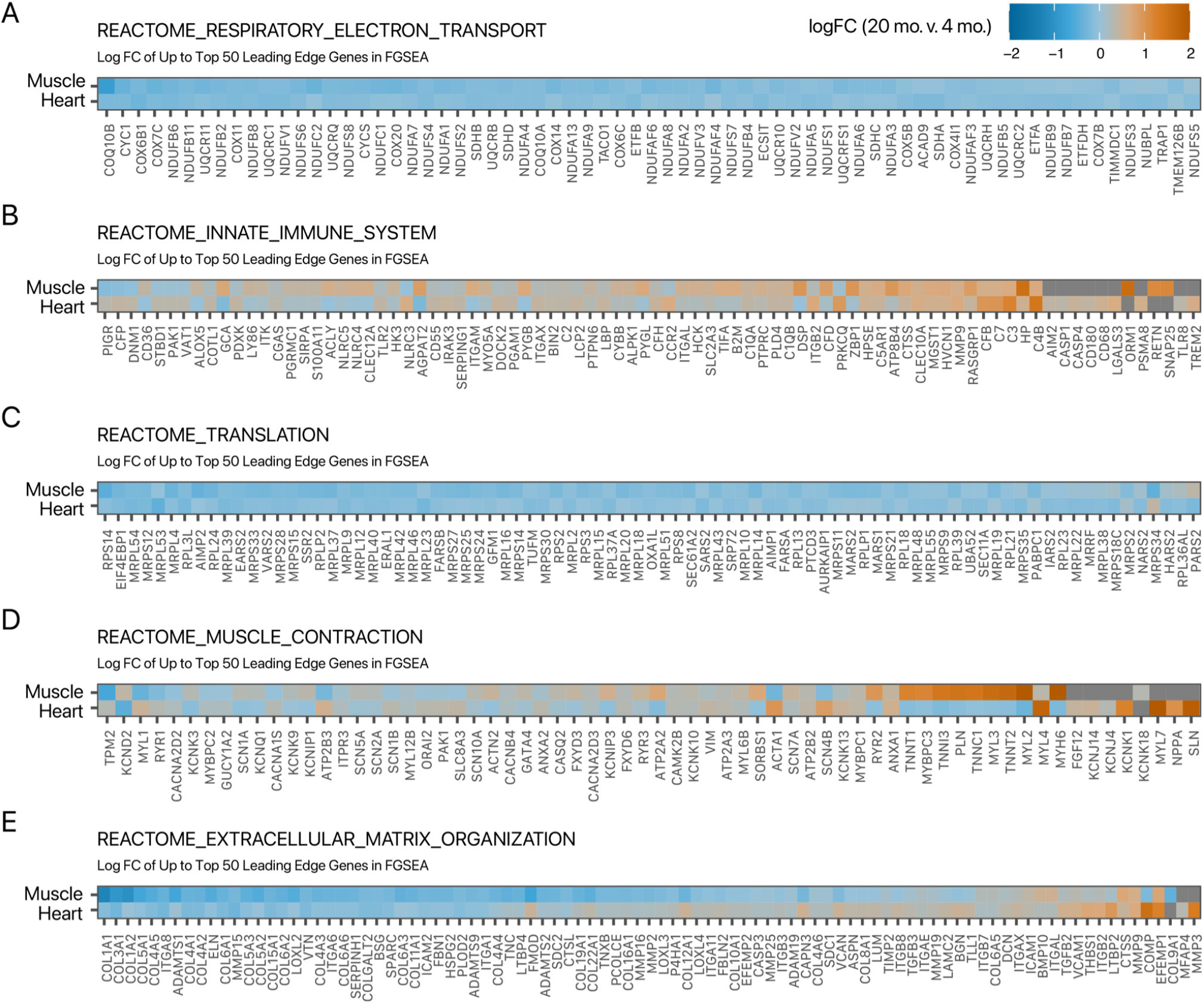
Heat maps of genes in selected enriched Reactome pathways. **A–E.** The heat maps show the log2 fold-changes (logFC) of aged vs. young adult mice in the heart and the skeletal muscle. Up to 50 leading edge genes responsible for FGSEA enrichment in each tissue are shown for **A.** respiratory electron transport; **B.** innate immune system; **C.** translation; **D.** muscle contraction; **E.** extracellular matrix organization. Fill color: logFC; missing values are in dark grey.

**Supplementary Figure S3.**
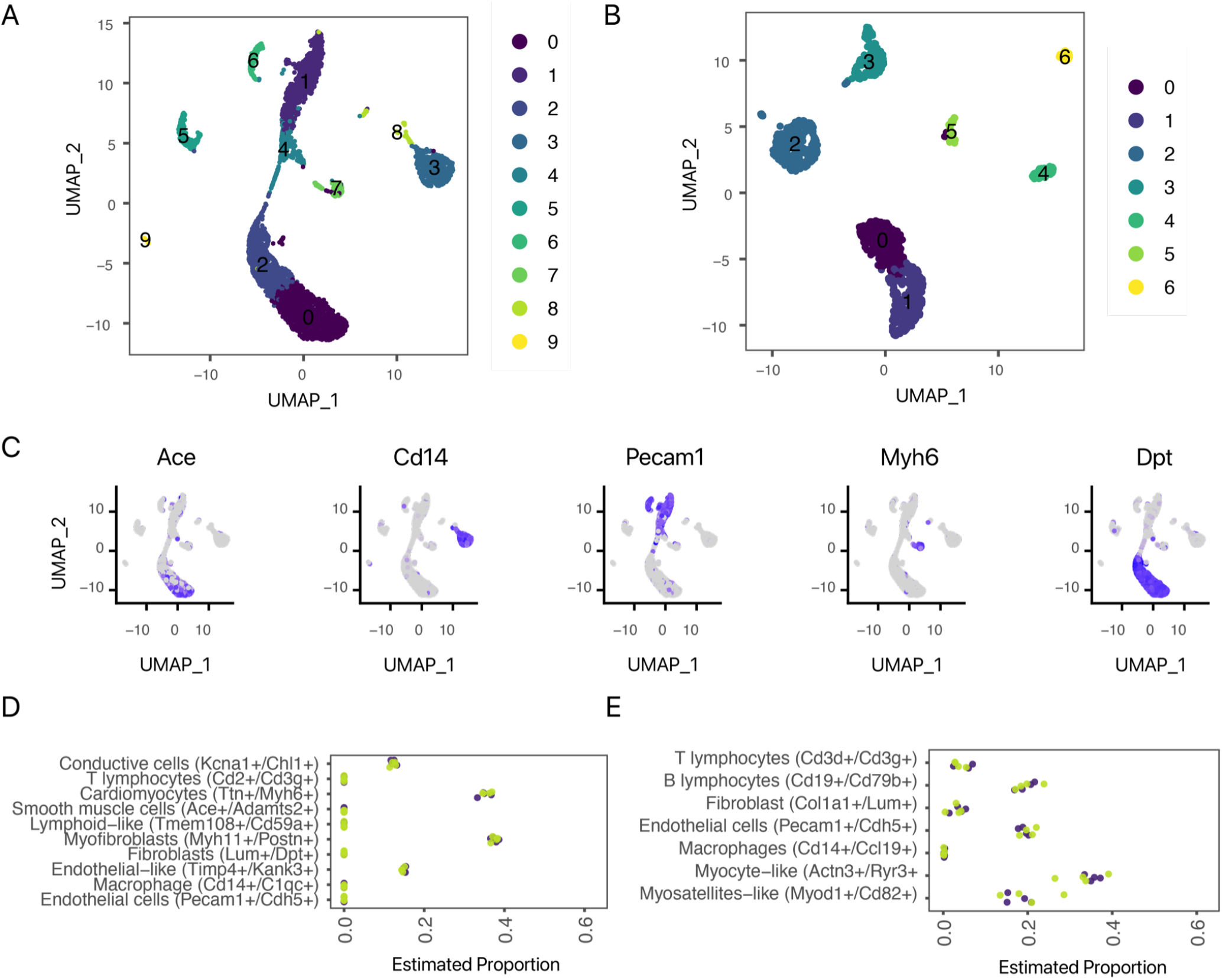
Estimation of cell type proportions. **A–B.** UMAP projection of Tabula Muris mouse (**A**) heart and (**B**) skeletal muscle FACS single-cell RNA sequencing data, with clusters found using the multilevel-refined Louvain algorithm implemented in Seurat. **C.** Tentative assignment of cell identity in the clusters was performed by manual interpretation of the top markers found by Seurat in each cluster, several examples of which are shown here. **D–E.** Estimated cell proportion from the bulk RNA-sequencing data in young adult (green) vs. early aging (purple) heart (**D**) and skeletal muscle (**E**).

**Supplementary Figure S4.**
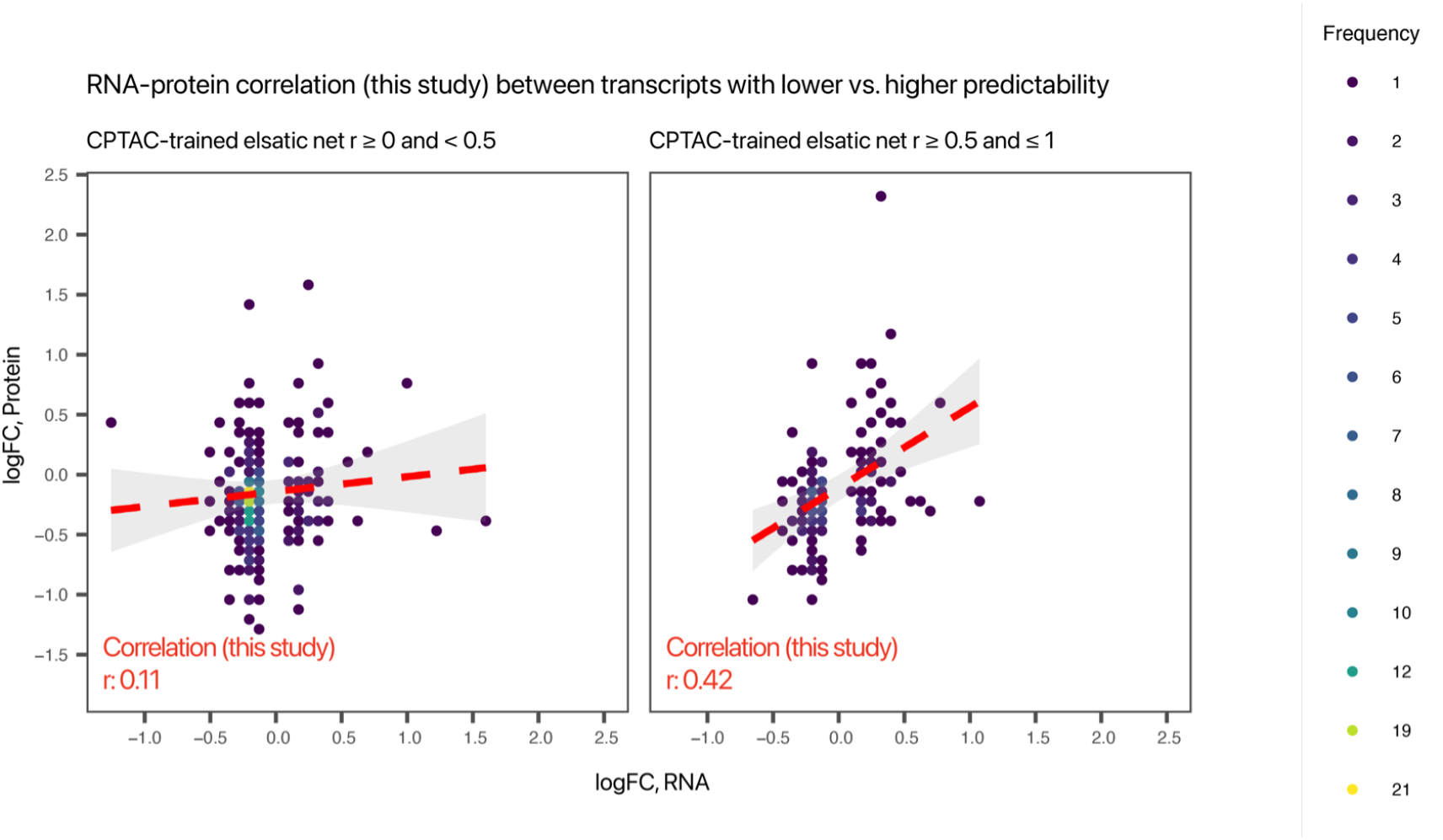
Transcripts with higher protein predictability show greater RNA-protein correlation in aging changes. Frequency scatter plot showing the correlation between RNA and protein level fold-changes in 20 months vs. 4 months in both tissues. X-axis: measured log2 fold change (logFC) at the transcript level in this study; y-axis: measured logFC at the protein level in this study. Left: transcripts with low predicted correlation with protein levels; right: transcripts with high predicted correlation with protein levels. Only transcript with nominal changes in this study (DESeq2 P ≤ 0.1) are included. Color: data point frequency within bin.

**Supplementary Figure S5.**
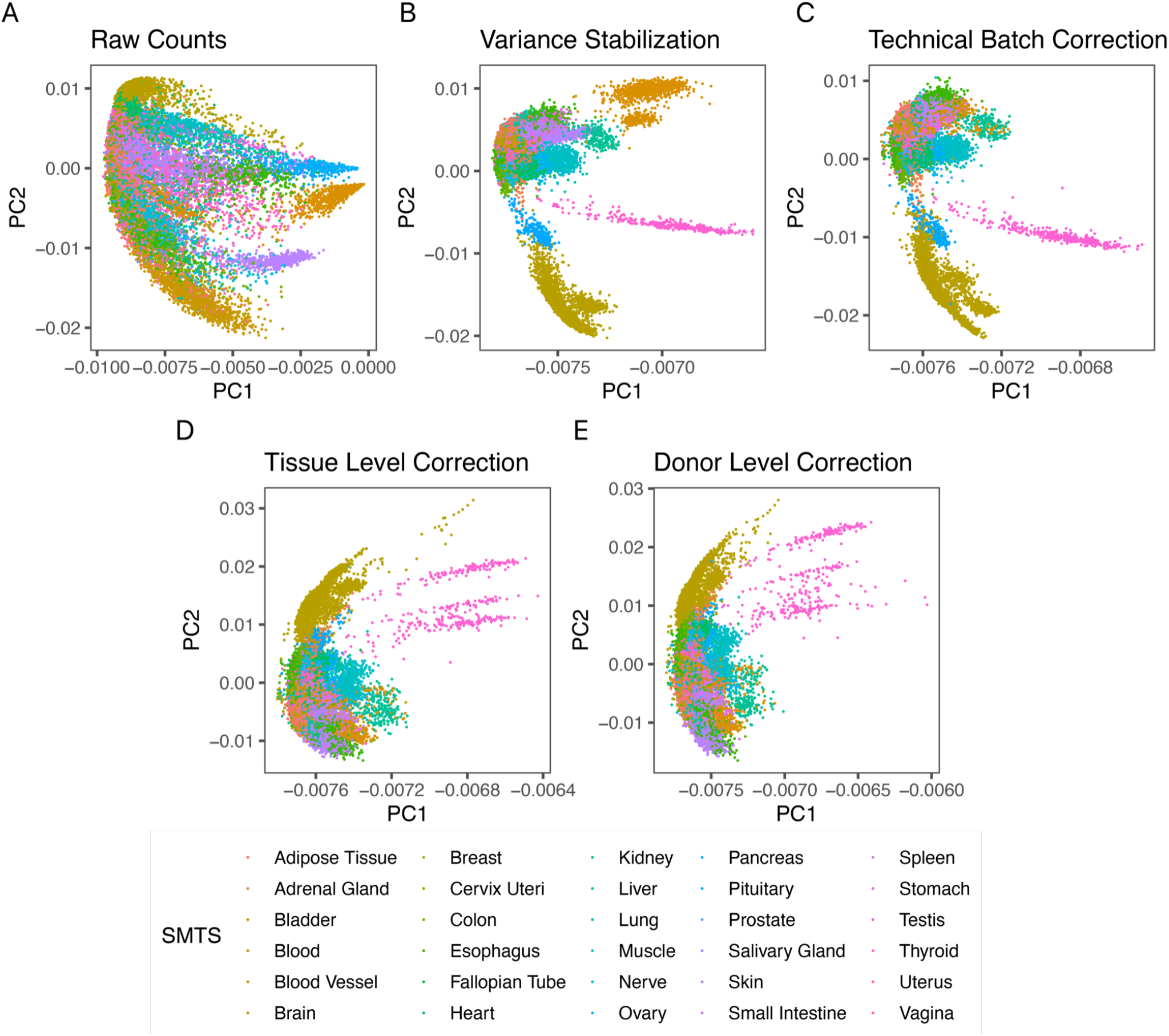
Normalization and batch correction of GTEx v8 data. From upper left to lower right, scatter plots showing the first two principal components of GTEx v8 transcriptome data: (**A**) raw counts; (**B**) variance stabilization transformed counts; (**C**) after ComBAT batch correction against two technical variables (extraction batch SMNABTCH and sequencing batch SMGEBTCH); (**D**) after ComBAT batch correction against tissue biological characteristics (ischemia time SMTSISCH); (**E**) after ComBAT batch correction against donor characteristics (death Hardy scale DTHHRDY). Each data point represents one tissue transcriptome. Color: GTEx v8 sample tissue metadata (SMTS).

**Supplementary Figure S6.**
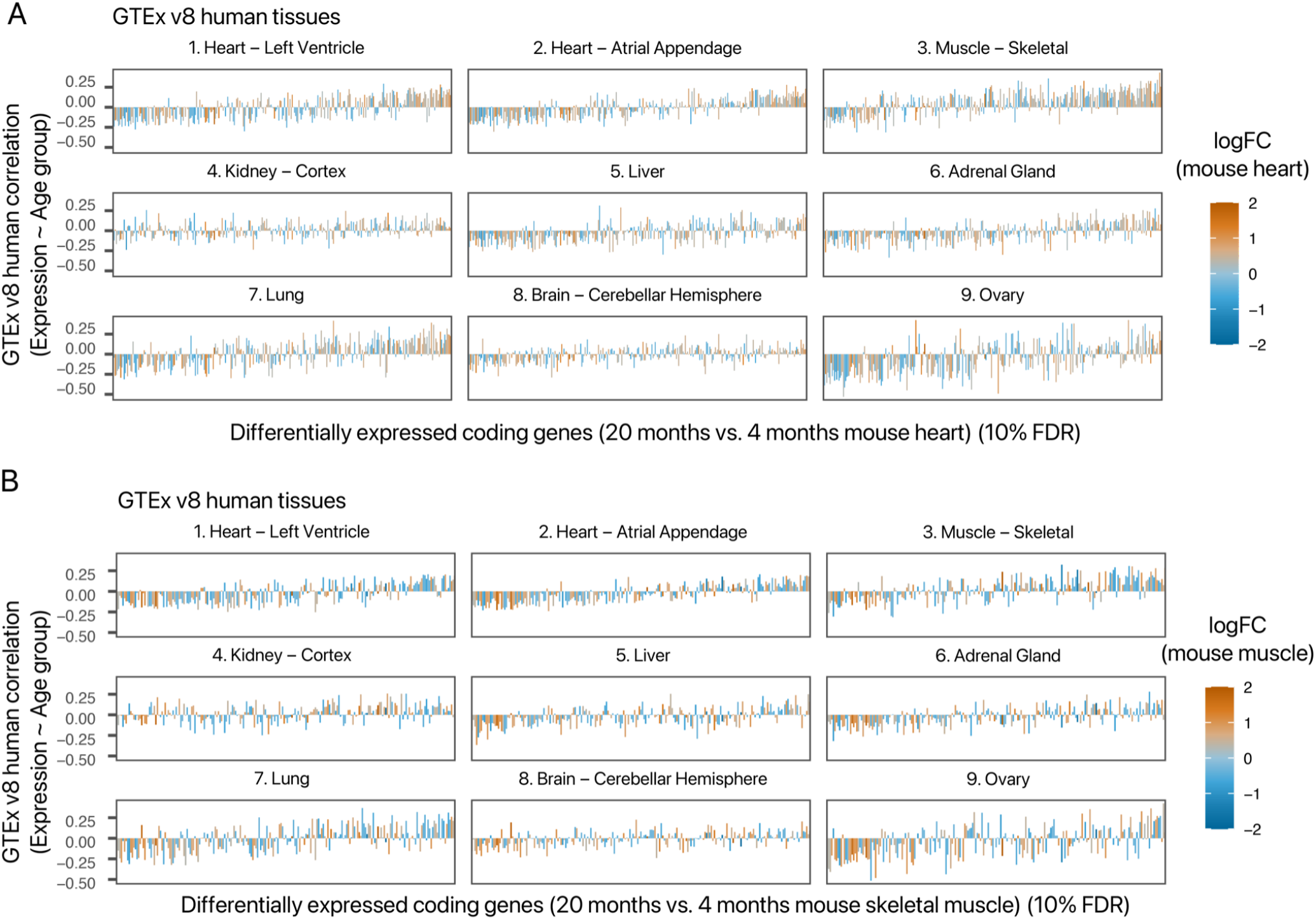
Age-expression relationships in human GTEx v8 data for the identified age-associated transcripts. Each bar chart shows the correlation coefficients between relative expression and age groups in 9 GTEx v8 human tissues for each of the differentially expressed genes in the mouse (**A**) heart and (**B**) skeletal muscle identified in this study. X-axis: differential transcripts in 20 months vs. 4 months mice; y-axis GTEx v8 correlation between expression and donor age group; fill-color: log2 transcript fold-change (logFC) in this study.

**Supplementary Table S1.** Animals used in the study. Dates of birth (DOB), dissection, heart weights, and tibia lengths are shown. LV: left ventricle; RV: right ventricle.

| ID | Dissection Date | Sex | DOB | Age (wks) | Body Mass (g) | Heart Mass (mg) | Atrium Mass (mg) | LV Mass (mg) | RV Mass (mg) | Tibia Length (mm) (3 repeated measures) | Avg. Tibia | HW/BW (mg/g) | LV/tibia (mg/mm) | RV/tibia (mg/mm) |
| --- | --- | --- | --- | --- | --- | --- | --- | --- | --- | --- | --- | --- | --- | --- |
| O12-F | 5/15/19 | F | 9/19/17 | 86 | 39.73 | 159.9 | 15.7 | 103.1 | 36.8 | 18.50 18.50 18.43 | 18.48 | 4.02 | 5.58 | 1.99 |
| O11-F | 5/15/19 | F | 9/19/17 | 86 | 33.39 | 127.3 | 9.8 | 85.4 | 34.9 | 18.64 18.60 18.60 | 18.61 | 3.81 | 4.59 | 1.88 |
| Y15-F | 5/9/19 | F | 1/15/19 | 16 | 21.29 | 103.9 | 6.9 | 72.8 | 20.6 | 17.83 17.77 17.75 | 17.78 | 4.88 | 4.09 | 1.16 |
| Y14-F | 5/9/19 | F | 1/15/19 | 16 | 22.17 | 107.4 | 6.1 | 75.7 | 23.9 | 18.01 17.95 17.98 | 17.98 | 4.84 | 4.21 | 1.33 |
| O14-M | 5/15/19 | M | 9/19/17 | 86 | 42.85 | 181.1 | 15.3 | 109.9 | 60.3 | 18.11 18.12 18.10 | 18.11 | 4.23 | 6.07 | 3.33 |
| O17-M | 5/16/19 | M | 9/19/17 | 86 | 37.31 | 150.9 | 9.0 | 104.7 | 34.8 | 18.33 18.32 18.29 | 18.31 | 4.04 | 5.72 | 1.90 |
| Y20-M | 5/14/19 | M | 1/15/19 | 17 | 31.75 | 127.6 | 7.8 | 95.2 | 24.5 | 18.15 18.10 18.11 | 18.12 | 4.02 | 5.25 | 1.35 |
| Y19-M | 5/14/19 | M | 1/15/19 | 17 | 30.53 | 135.8 | 10.2 | 94.7 | 31.1 | 18.16 18.14 18.17 | 18.16 | 4.44 | 5.22 | 1.71 |

## Supplementary Data

**Supplementary Data S1** (.txt): DESeq2 output for all transcript comparisons (20 months vs. 4 months) in the heart.

**Supplementary Data S2** (.txt): DESeq2 output for all transcript comparisons (20 months vs. 4 months) in the skeletal muscle.

**Supplementary Data S3** (.txt): Correlation coefficients and R^2^ between transcript-predicted and empirical protein level data for 11,896 transcripts in CPTAC samples in the trained model.

**Supplementary Data S4** (.xlsx): Enriched Reactome terms among transcripts that predict proteins poorly (r ≤ 0.3) or relatively well (r ≥ 0.5)

**Supplementary Data S5** (.xlsx): Ionquant output label-free quantitative data of proteins in the heart and the skeletal muscle

**Supplementary Data S6** (.txt): DESeq2 output for all transcript comparisons (20 months vs. 4 months) in the kidney.

